# Identifying Gene-wise Differences in Latent Space Projections Across Cell Types and Species in Single Cell Data using scProject

**DOI:** 10.1101/2021.08.25.457650

**Authors:** Asher Baraban, Brian S. Clark, Jared Slosberg, Elana J. Fertig, Loyal A. Goff, Genevieve Stein-O’Brien

## Abstract

Latent space techniques have emerged as powerful tools to identify genes and gene sets responsible for cell-type and species-specific differences in single-cell data. Transfer learning methods can compare learned latent spaces across biological systems. However, the robustness that comes from leveraging information across multiple genes in transfer learning is often attained at the sacrifice of gene-wise precision. Thus, methods are needed to identify genes, defined as important within a particular latent space, that significantly differ between contexts. To address this challenge, we have developed a new framework, scProject, and a new metric, projectionDrivers, to quantitatively examine latent space usage across single-cell experimental systems while concurrently extracting the genes driving the differential usage of the latent space between defined contrasts. Here, we demonstrate the efficacy, utility, and scalability of scProject with projectionDrivers and provide experimental validation for predicted species-specific differences between the developing mouse and human retina.

## MAIN

The growing size and complexity of scRNA-seq datasets has necessitated dimensionality reduction techniques including PCA, CCA, ICA, NMF, CoGAPS, UMAP, TSNE, scVI, and scARCHES to extract biologically meaningful latent spaces^1–61,3,7–10^. These learned latent spaces serve as representations of cell type, cell state, or distinct biological processes which exhibit coordinated regulation, such as induction of gene expression following pathway activation. Transfer learning (TL) is a powerful tool for comparing latent space usage across single-cell studies^8,10–16^.

TL methods can overcome batch effects and other unwanted noise by conditioning on the shared structure of conserved biological processes. Using latent spaces defined in a source dataset, TL methods rapidly identify shared biological processes in a new target dataset^10,12,17^. By facilitating post hoc analysis of learned latent spaces, unsupervised dimension reduction techniques are often used for information discovery within single-cell technologies^3^. These techniques overcome many of the technical challenges of single-cell data including gene dropout and sparsity^1,3,4,12^. Furthermore, latent spaces have been used in a variety of transfer learning contexts to quickly and robustly annotate data across different experimental conditions or biological systems^10–12,17–19^.

Within biological research, model organisms are pivotal to studies seeking to understand the biological mechanisms central to human development and disease, prefaced on understanding evolutionarily conserved biological features to advance our knowledge of human biology. For example, key biomarkers and/or pathways are often evolutionarily conserved allowing investigators to use model organisms to discover and study their function^20,21^. Conversely, adaptations that confer desirable functionality can serve as the inspiration for therapies, e.g. the ability of zebrafish to regenerate their retina following injury^22^. Thus, the ability to find genes and gene sets involved in key celltype and species-specific differences can yield important biological insights.

However, within scRNA-seq studies, both batch effects and the inherent loss of resolution in dimensionality reduction makes it difficult to derive gene level biological insights across latent spaces^8,23,24^. These differences are especially important for cross-species analysis where the evolutionary and functional relationship between cells and/ or genes responsible for a conserved biological process are often unknown^21,24^. The robustness that comes from leveraging information across multiple genes in transfer learning systems is attained at the sacrifice of gene-wise precision^3,23,25^. Thus, methods are needed to identify the underlying genes driving meaningful differences between contrasts of interest derived from latent space projections. To address this challenge, we have developed a new framework, scProject with projectionDrivers, to quantitatively examine latent space usage across single-cell experimental systems while concurrently extracting the genes driving the differential usage of the latent space across the defined testing parameters. scProject implements an array of TL methods by extending the widely used python scRNA-seq workflow provided by ScanPy^26^. With scProject, any provided latent space can be leveraged to extract gene-wise insights from other datasets within an AnnData-backed ScanPy genomics pipeline.

Additionally, we developed the projectionDrivers metric to identify genes driving the projection of a latent space in a target dataset to understand the specific differences in latent space usage, and therefore biological process differences, between two conditions. We then apply this approach to identify gene-level differences in a clinically relevant biological system, the developing vertebrate retina. As the mouse developing retina is a highly tractable experimental system displaying numerous conserved evolutionary features across vertebrates, many retinal studies utilize mice to facilitate studies on human retinal development and disease. More than 10% of American adults over 40 suffer from visual impairments that affect the retina, including blindness, age-related macular degeneration, diabetic retinopathy and glaucoma (National Eye Institute - NEI). Current projections estimate that the number of cases of visual impairment will double over the next 40 years (NEI). Here, we demonstrate scProject’s efficacy and utility for inter-tissue and cross-species analyses using previously published sets of latent spaces from the developing mouse and human retina^24,27^. Using scProject, we highlight previously undescribed gene expression differences between mouse and human retinal development, identifying and providing independent experimental validation of evolutionarily divergent expression of genes implicated in human retinal disease. Finally, we showcase scProject’s scalability by analyzing a dataset comprising four million cells.

## RESULTS

### Overview of the scProject workflow and projectionDrivers

We have developed a scalable framework, scProject, for the projection of target datasets into previously defined latent spaces. Given the widespread adoption of dimension reduction techniques for single-cell analysis, scProject starts with a user-provided set of previously learned latent spaces. Each space is encoded in a single vector of gene weights, the structure of which defines the co-expression of genes associated with a given biological or technical feature of interest. These spaces can define both discrete (i.e. cell type) and continuous (i.e. cell cycle position) features learned from a source dataset. Together, a set of latent spaces forms a basis defining the experimental system from which it was learned.

At its core, scProject performs a change of basis on a new independent target dataset into the basis defined by the source dataset—in essence mapping, or projecting, the new samples into the latent space defined by the source data. This change of basis quantifies the usage of each individual latent space in the target dataset, resulting in a new matrix (pattern matrix) which encodes this information in the form of coefficients assigned to the latent space (basis vector) for each sample/cell. To quantify the usage of the latent spaces in the target dataset, scProject performs a non-negative elastic net regression using scikit-learn^28^.

We chose to constrain solutions to nonnegative values to reflect true biological constraints; however, scProject allows users to use unconstrained elastic net regression allowing for the use of latent spaces containing negative weights such as those learned by PCA. Negative gene expression does not occur in nature, and analogous constraints have been previously demonstrated to improve biological inference^1,6,25,29^. After the regression, positive coefficients indicate usage of a latent space in a target sample. Thus, biological processes that are shared across datasets will result in positive values, while dataset-specific factors such as dataset-specific technical artifacts will be minimized^11–13^. The inherent conservation of biological processes across systems, and subsequently datasets, makes transfer learning a valuable tool for cross-species comparisons^12–14^.

To amplify signal and discard spurious latent space usage during the projection step, we offer additional tools within scProject. The elastic net regression in scProject both encourages sparsity, a known feature of single-cell data, while also handling the potential for collinearity (i.e. linearly dependent latent spaces) since biological processes are often interdependent. scProject allows the user to choose the weight of regularization and the percent of lasso regression. Stronger regularization forces the model to pick the most representative latent space thereby also serving as an informal measure of robustness of latent space usage for a cell type or cluster. Lower regularization allows for the identification of more general relationships between latent space usage and existing annotations, but may also result in more spurious associations. Users can assess the sensitivity of latent space usage to regularization parameters thereby testing the overall robustness of a latent space’s usage, for which we provide an example in the context of finding the latent spaces most consistent with a cell type of interest (Supplemental notebook Microglia.ipynb).

As visualization of the target data projected into the latent spaces can quickly summarize trends and help prioritize latent spaces of interest, we developed several plot based assessment methods. The first visualization uses UMAP^30^ to reduce the dimensionality of the target data’s projection in the source latent spaces to two dimensions. Samples can be colored by their annotation or coefficient of usage for a specific latent space. When combined with known annotations such as cell type, this global representation is useful to illustrate the contextual use of a specific latent space, and can often provide insight into the biological process it may represent in the target dataset.

The second visualization method uses Pearson correlation to assess the strength of the relationships between each latent space and any annotated features. The resulting heatmap of coefficients highlight latent spaces strongly associated with particular aspects of the new data, e.g. a specific single-cell type or biological process shared across diseased, but not control samples.

To identify genes of interest within a latent space, we provide several methods to interrogate the gene weights and individual gene contributions to a projection analysis. The ‘importantGenes’ function lists the genes in a latent space expressed above a provided threshold. The ‘geneSelectivity’ function measures the selectivity of a gene’s expression in a given latent space and displays a bar plot illustrating usage of the gene throughout the latent spaces.

Finally, the projectionDrivers tool can be used to identify genes driving differential latent space usage between two conditions within a target dataset. Importantly, the nature of projectionDrivers is such that it provides a pairwise comparison between user-defined conditions, and the definition of the comparison conditions can be adjusted to interrogate different aspects of a latent space usage. For example, discretization of a target dataset into cells that ‘use’ a particular latent space and cells that do not (Supplemental notebook 39ExpressVs.ControlRodsVs.BipolarCells.ipynb), provides an intuitive grouping to examine the specific genes involved in the projection of the latent space globally across the target dataset.

Additionally, a comparison of two annotated cell types, both of which demonstrate significant latent space usage in the target dataset, can be used to explore the cell-type-specific differences in latent space usage (Supplemental notebook 39ExpressVs.ControlRodsVs.BipolarCells.ipynb). Importantly, the use of latent spaces examines the dataset as a whole, outside of singular presence or absence of gene expression - a vitally important aspect given the sparsity and drop-out rate of scRNA-seq data. However, the underlying statistics contained here provide gene-level resolution to determine individual gene contribution to the biological process in question.

### Identifying gene-wise drivers of a latent space projection

While the coregulation of genes responsible for a specific biological process is often conserved across species or cell types, there may be subtle differences in gene use that contain important biological information. For example, certain subsets of a signaling pathway may be differentially used across cellular states, cell types, ages, or species. Most approaches to transfer learning-based representation of biological processes generally do not account for subtle differences in latent space use or individual gene expression differences across conditions.

However, divergence of biological processes, including the gene sets associated with a particular process, may be of interest. To address this need, we have defined the ‘projectionDrivers’ function which determines if the differential gene expression between two subsets of cells (clusters as e.g.) is driving latent space usage. Specifically, projectionDrivers adapts the previously described Bonferroni correction for confidence intervals^31^ to select for these genes.

scProject provides statistical functions to allow a user to rigorously find genes driving the differential usage of a latent space between two subsets of cells within a target dataset. We foresee these cellular subsets being defined in many ways: their latent space use or lack thereof, cell type annotations, marker gene expression, or other user-defined annotation parameters.

To determine the significance of differential gene expression between two conditions of interest, the ‘BonferroniCorrectedDifferenceMeans’ function outputs confidence intervals for genes that are potentially responsible for the difference in usage of the queried specific latent space between the two populations at a specified Bonferroni corrected level of significance. Through utilization of a log transformation prior to implementing the BonferroniCorrectedDifferenceMeans function, we produce confidence intervals around the log_2_ fold change of each gene. Under these conditions, the log_2_ fold change is an unbiased estimator.

projectionDrivers takes advantage of these facts to correct for batch effects by taking the difference of the mean of the log transformed expression values within the same batch. We can then compare the log_2_ fold change across batches and therefore species to make accurate, unbiased comparisons. While most algorithms use a joint learning paradigm to align multiple datasets,^14,32,33^ we assume each dataset is from a different batch, and the batch effect is modeled as every measurement of a given gene within a batch is multiplied by a constant factor. This ratio improves upon previous joint learning algorithms by correcting for unmatched batch effects.

Mathematically, projectionDrivers constructs two sets of confidence intervals for each gene around the difference in mean gene expression values for two different subsets of cells in a given single-cell dataset (e.g. CI of log fold change difference between two annotated cell types; Supplemental Methods). The first is a standard Bonferroni corrected confidence interval on the log_2_ fold change of the mean gene expression estimates for cells in two subsets of cells. A greater than 99% confidence interval that does not overlap with zero suggests that the gene is significantly differentially expressed between the two populations. By log-transforming before input, the function creates confidence intervals around the log fold change in mean gene expression between the two clusters thereby removing batch effects. The Bonferroni corrected confidence interval, 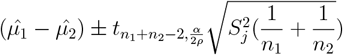, where 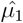 and 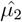 are the vectors of the mean of log transformed gene expression for subset 1 and 2 respectively. *n*_1_ and *n*_2_ are the number of cells in subset 1 and 2 respectively. *α* is the level of confidence, 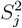 is the sample variance of gene *j*, and *ρ* is the number of genes.

The second set of confidence intervals are derived as a weighted Bonferroni corrected confidence interval, where we weight the difference in mean gene expression by the latent space weight for each given gene. This second confidence interval ensures that the gene has a high enough weight in the latent space. In other words, by taking the product of the mean difference and the weight of the gene in the latent space, we ensure that the gene is both used in the latent space and differentially expressed. We define the second weighted confidence interval as 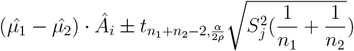 where *Â*_*i*_ is the *i*th latent space with an L1 norm equal to the number of nonzero entries in the latent space.

The projectionDriver genes are defined as those in which both confidence intervals do not include 0. These conditions identify genes that are both differentially expressed between the two subpopulations of interest and significantly used in the latent space. Therefore, these genes are driving the differential use of the latent space. In this way, we disentangle the genes aggregated by dimensionality reduction techniques. Our projectionDriver approach can clarify the outputs of dimensionality reduction-based latent space models by isolating genes of interest from the hundreds or thousands of genes expressed in a latent space.

### Application to latent spaces from retina development

To illustrate the ability of a set of latent species to make cell type predictions across species, we used a set of 80 latent spaces learned on 120,804 single-cells from whole mouse retinas at 10 select developmental time points, ranging from prior to the onset of neurogenesis (E11) through terminal fate specification (P14)^27^. These 80 latent spaces spanning mouse retina development capture both discrete (e.g. cell types) and continuous (e.g. cell cycle) biological processes ^27^ and have been successfully used for cross-species TL^12,24^.

We evaluated the ability of these mouse latent spaces to recover known cell type annotations from a comparable single-cell RNA-Seq dataset from the developing human retina^24^. The human dataset is comprised of 118,555 single-cell transcriptional profiles from whole developing retinas obtained at 9, 11, 12, 13, 14, 15, 16, 17, 18, 19, 22, 24, and 27 gestational weeks (GW), and macular and peripheral samples from 20 GW and 8 days postnatal (PND). Biological and technical replicates were performed at GW24 and GW19, respectively. Samples were profiled to a mean depth of 3,472.61 unique molecular identifiers (UMIs; SD = 1,862.78) and 1,502.14 genes (SD = 589.41) per cell^24^.

We projected both the original mouse and human log-transformed gene expression values into the aforementioned mouse latent space using scProject. From the projection, each cell has a weight associated with each individual latent space. We used thirty percent of the cells to train 80 random forest classifiers to predict cell type, one for each latent space. We measured the predictive power of each classifier (latent space) by measuring the percentage of each cell type they predicted correctly on the remaining seventy percent of the data.

We then used AUC scores after the lowLasso adjustment to identify latent spaces associated with high predictive power for a given cell type. Using an AUC threshold of 0.80, we identified 9 out of 80 latent spaces with significant predictive power for one or more cell types in both the mouse and human dataset (Fig. 1a). 4 of these 9 latent spaces were predictive of a single-cell type each, and these predictions were consistent across both species. The remaining 5 had high predictive power for no more than 2 developmentally-related cell types each in either species, suggesting that these predictions may arise from developmentally related cell types.

**Figure 1:**
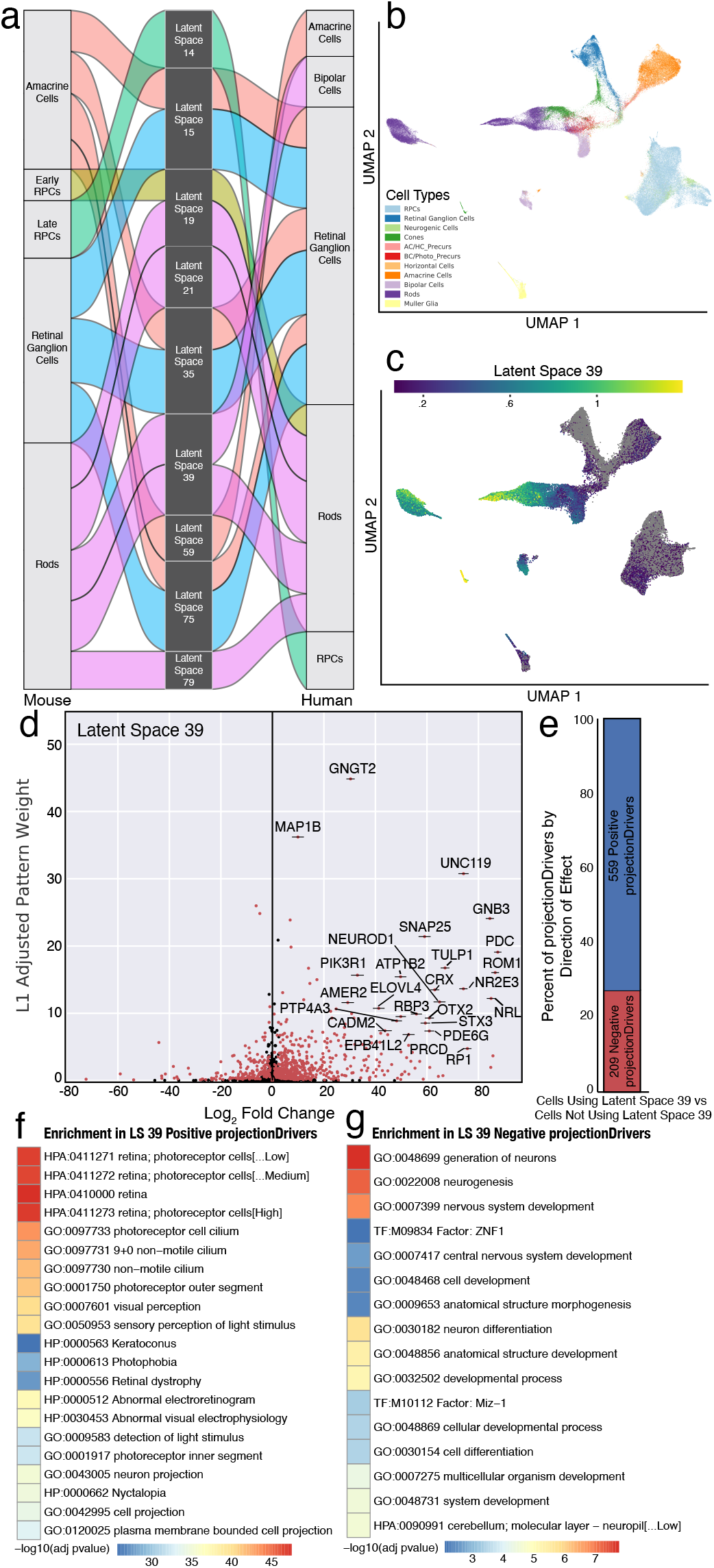
Identification of projectionDriver genes for a single latent space projection between developing mouse and human retina. a) Alluvial plot illustrating the predictive power of latent spaces across human and mouse developing retina. The width of the connection from cell type to latent space is proportional to the percentage of cells predicted correctly by a random forest trained on that latent space. Only latent spaces that had an AUC of ROC of at least .8 are included. b) UMAP of developing human retinal cells based on the projected mouse pattern matrix projected annotated by cell type. c) UMAP coordinate plot colored by the coefficient of latent space 39 highlighting the cells using latent space 39. d) Volcano plot of log_2_ fold change vs L1-adjusted latent space weight. Each point represents a gene. Those colored in red are projectionDrivers and those colored in black are not. The annotated genes are the top ranked genes by product of log_2_ fold change and L1-adjusted latent space weight. e) Bar plot showing the proportion of projectionDriver genes identified as either positively associated with latent space 39 (blue) or negatively associated with latent space 39 (red). f-g) Significant gene set enrichment ontologies for positive and negative projectionDrivers.

### Projection drivers of mouse latent space 39 across the developing human retina

To illustrate the application of projectionDrivers to identify genes associated with a particular latent space projection, we next focused on latent space 39. In the source mouse dataset, latent space 39 has high predictive power for rod photoreceptors (Fig. 1a). After projecting the human retina dataset (Fig. 1b) we identified 25,346 cells with significant projections (defined as coefficients greater than .5) into latent space 39 (Fig. 1c).

To identify the genes driving the usage of latent space 39 across all cells in the human dataset regardless of cell type, we set up a projectionDrivers comparison between those cells with a coefficient of the latent space above .5 and those not using the latent space at all (i.e. zero coefficient of latent space 39). This contrast identified 559 genes as positive projection drivers associated with the use of latent space 39 in the developing human retina (Fig. 1d-e). Conversely, we also identify 209 genes with high weight in latent space 39 that were enriched in cells with zero projection weights for latent space 39, suggesting that these genes are not involved in the latent space 39 usage in the developing human retina in the same manner as they are in the mouse dataset. To illustrate the relationship between predicted gene involvement in latent space 39 (gene weight) and the relative difference in gene expression between cells with a positive projection vs those without in the target human dataset, we created a volcano plot (Fig. 1d) and a ranked CI plot (Supplemental Fig. 1) to visualize the genes driving the usage of latent space 39 in the dataset. The majority of the projectionDriver genes are enriched in the cells with high-scoring projections in latent space 39. Gene set enrichment analysis (GSEA) of the 559 positive projectionDriver genes indicated that this set is enriched for genes associated with photoreception, visual perception, detection of light, and other relevant ontologies for a latent space associated with rod photoreceptors (Fig. 1f-g).

### Projection drivers of mouse latent space 39 between human rods and bipolar cells

While latent space 39 is predictive of rod photoreceptors in mice, we identified mouse latent space 39 as predictive of both rods and bipolar cells in the developing human retina dataset (Fig. 2a). To better understand how latent space 39 is used differently across the two cell types, and to identify specific genes that may be differentially contributing to the significant latent space usage with each cell type, we performed a projectionDrivers analysis of the log_2_ fold change of the mean gene expression levels between rods and bipolar cells in the target human dataset.

**Figure 2:**
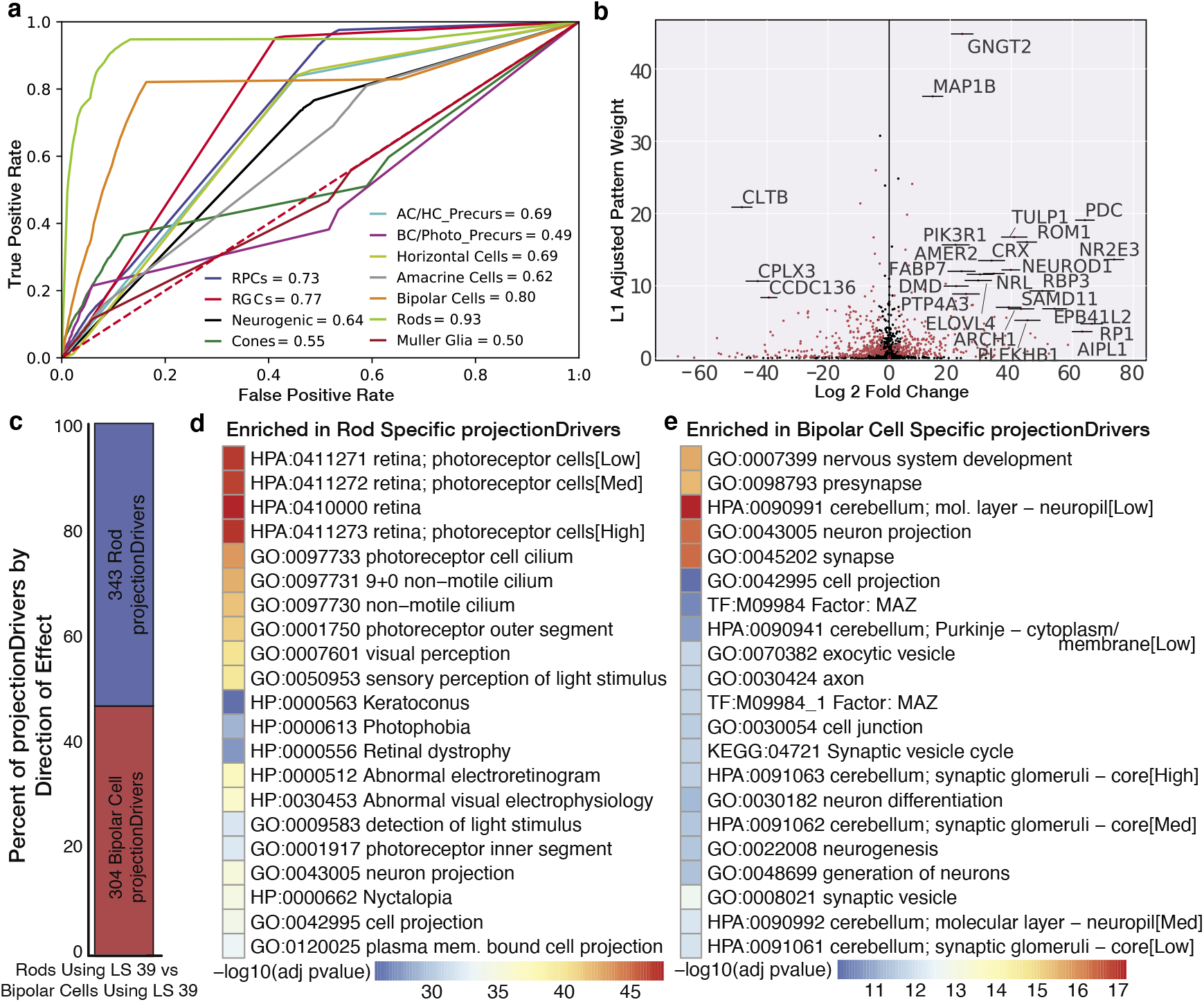
Analysis of Rods versus Bipolar Cells in Mouse Latent Space 39. a) ROC curves for each cell type from a random forest trained on latent space 39. b) Volcano plot with log_2_ fold change in gene expression between rods and bipolar cells vs L1-adjusted latent space weights. Each point represents a gene. Those colored in red are projection-Drivers and those colored in black are not. The annotated genes are the top ranked genes by product of log_2_ fold change and L1-adjusted latent space weight. c) Bar plot showing the proportion of projectionDriver genes identified as either bipolar projectionDrivers (red) or rod projectionDrivers (blue) d-e) Significant gene set enrichment ontologies for rod and bipolar cell specific projectionDrivers.

We visualized the confidence intervals from projectionDrivers with a volcano plot with log_2_ fold change between the two cell types on the abscissa and L1 norm adjusted gene weights on the ordinate axis (Fig. 2b). The genes annotated in the right quadrant are enriched in rods and have high weights in latent space 39. Similarly, the genes annotated in the second quadrant are enriched in bipolar cells and have high weights in latent space 39 (Fig. 2b). This contrast identified 343 genes as significant projection drivers for latent space 39 use in rods, and 304 genes as significant projection drivers for the same latent space 39 in bipolar cells (Fig. 2c). Gene set enrichment analysis of projectionDrivers for the rod cell type was significant for biological processes and diseases specific to photoreceptors and light detection by contrast gene set enrichment analysis of projectionDrivers for the bipolar cells was significant for biological processes related to neuronal development and function (Fig. 2d-e).

### ProjectionDrivers identifies cross species differences between developing mouse and human retina

As numerous retinal diseases result from dysfunction of the light sensing photoreceptor cells of the retina, we chose to apply scProject to identify differences in photoreceptor development and specification between mouse and human. Since scProject takes advantage of the batch correction inherent in ratio comparisons, we chose to evaluate the weighted difference in expression between photoreceptors and all other retinal cells (Rods+Cones vs All) within both mouse and human.

The first challenge is mapping genes across species to facilitate direct comparison of their expression. scProject provides a function ‘orthologMapper’, which takes as input a csv file from biomart^34^ which contains lists of orthologous genes between mouse and human and adds an annotation to the dataset of the orthologous genes. After generating the ortholog mapping, we then projected both the mouse and human datasets onto the same set of latent spaces learned from time course scRNA-seq data of the developing human retina^24^. This projection identified latent space 75 as being used across rods and cones in both species (Supplemental Fig. 2a-b). As expected, many of the highly weighted genes in latent space 75 play essential roles in the phototransduction pathway for rod or cone photoreceptors (*AIPL, RCVRN, GNB3*^*35–37*^) or photoreceptor development (*RXRG, PRDM1, NEUROD1*^*38–40*^).

We next sought to determine the extent to which the genes that comprise latent space 75 are differentially expressed across the two species. Using projectionDrivers, we first determined the confidence intervals of the log fold change difference between the annotated photoreceptors (rods and cones) and the remainder of the cell types in both human and mouse. We then compared the confidence intervals between species, ranking the genes by distance between the confidence intervals divided by the average length of the confidence interval multiplied by their weight in the latent space. In this way, projectionDrivers generated a prioritized list of high confidence genes, meaning those that are both highly weighted in the latent space and differentially expressed (Supplemental Fig. 2c).

Thus, genes identified by projectionDrivers are responsible for key species specific differences in the otherwise conserved biological process of phototransduction and photoreceptor development. For example, *KCNV2* and *RPGRIP1* display near binary differential usage across the species. *KCNV2*, a gene implicated in cone-rod dystrophy^41^, displays high expression within human rods and cones, but is only minimally detected within the mouse scRNA-seq dataset (Fig. 3a-c). Conversely, *RPGRIP1*, a protein required for normal photoreceptor disk morphogenesis and mutated in forms of Leber’s Congenital Amaurosis^42^, was only weakly detected in the human dataset, but displays robust expression in mouse rod and cone photoreceptors (Fig. 3d).

**Figure 3:**
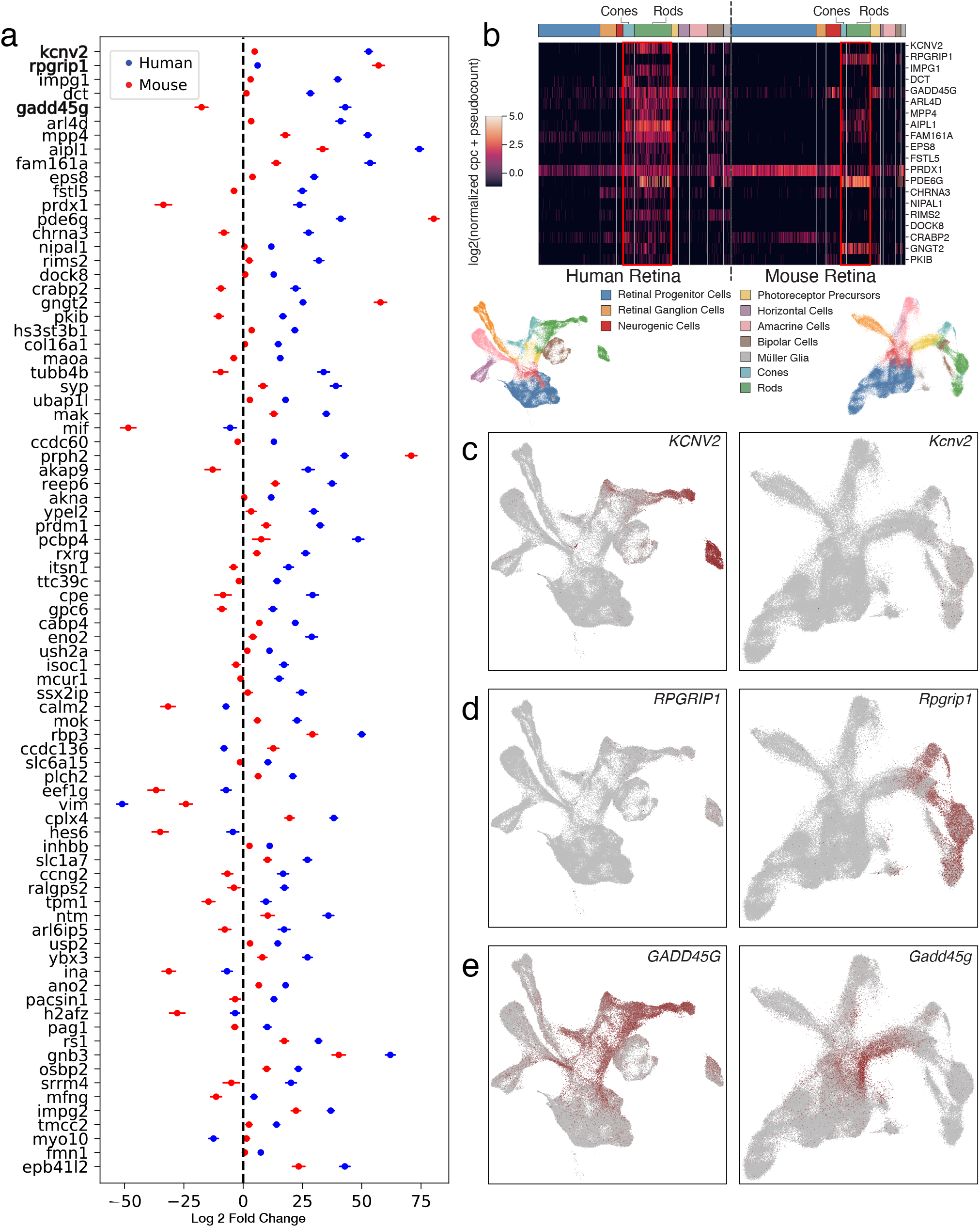
Human and mouse comparison of latent space 75 usage in rods and cones versus all other retinal cell types. a) Each point represents the log_2_ fold change of mean gene expression between rods and cones in human or mouse. Error bars on the points represent the Bonferroni corrected confidence intervals. Genes are ranked by distance between means divided by the average length of the confidence interval. b) Heatmap showing the log_2_ normalized gene expression across species and cell type for the top projectionDriver genes from the latent space 75 analysis of photoreceptors vs all other retinal cell types c-e) UMAP showing differential expression of selected genes between mouse and human developing retina. Each point represents data obtained from a single cell. The color of each point represents the expression level of that gene.

The true power of projectionDrivers can be seen in the more nuanced differences in *GADD45G/Gadd45g* expression (Fig. 3e). While both species use this gene extensively, *Gadd45g* is utilized in multiple latent spaces which correspond to different biological processes. For example, in the developing mouse retina, *Gadd45g* expression is enriched within neurogenic retinal progenitor cells (RPCs) and immature neurons (Fig. 3e, right panel), but its expression is reduced as cells mature. In contrast, in the developing human retina, *GADD45G* expression is maintained in photoreceptor cells into adulthood (Fig. 3e, left panel). This previously undescribed biological divergence was confirmed in two independent scRNA-seq datasets profiling transcript expression in adult human and mouse retina^43,44^, highlighting expression of *GADD45G* expression within human, but not mouse, rod photoreceptors (Supplemental Fig. 3a, 3b) and demonstrating the ability of projectionDrivers to identify gene drivers of species specific differential latent space usage.

We next expanded this projectionDrivers analysis to include scRNA-seq datasets from two additional model systems: human ES-cell/iPS-cell derived retinal organoids and the macaque retina^45,46^. The retinal organoid scRNA-Seq dataset represents an *in vitro* system mimicking *in vivo* human retinal development. The macaque retinal single-cell dataset represents a model system that highlights biology conserved between human and a closely related primate species. An additional advantage of the macaque as a model system for the human retina is that they share the fovea, a high acuity vision center, a feature absent from the *in vitro* retinal organoid system and mouse model.

To expand upon our previous analysis, we generated 4 sets of confidence intervals using projectionDrivers(Rods + Cones vs. All) weighted by feature 75. We then created a ranking which prioritized differential log fold change of the genes identified by projectionDrivers among all 4 species (Fig. 4). A highly ranked gene from this analysis is Dopachrome tautomerase (*DCT*; Fig. 4, 5). DCT, also known as tyrosinase-related protein 2 (TYRP2), functions as an enzyme in the melanin synthesis pathway^47^, converting DOPA-chrome to 5,6-Dihydroxyindole-2-carboxylic acid during the conversion of tyrosine to Eu-Melanin. This analysis revealed that DCT is highly expressed in human photoreceptors, particularly cones. Earlier studies had shown prominent DCT expression in retinal pigment epithelium (RPE) in both human and mouse^48,49^; however, its selective expression in human cones has not previously been characterized.

**Figure 4:**
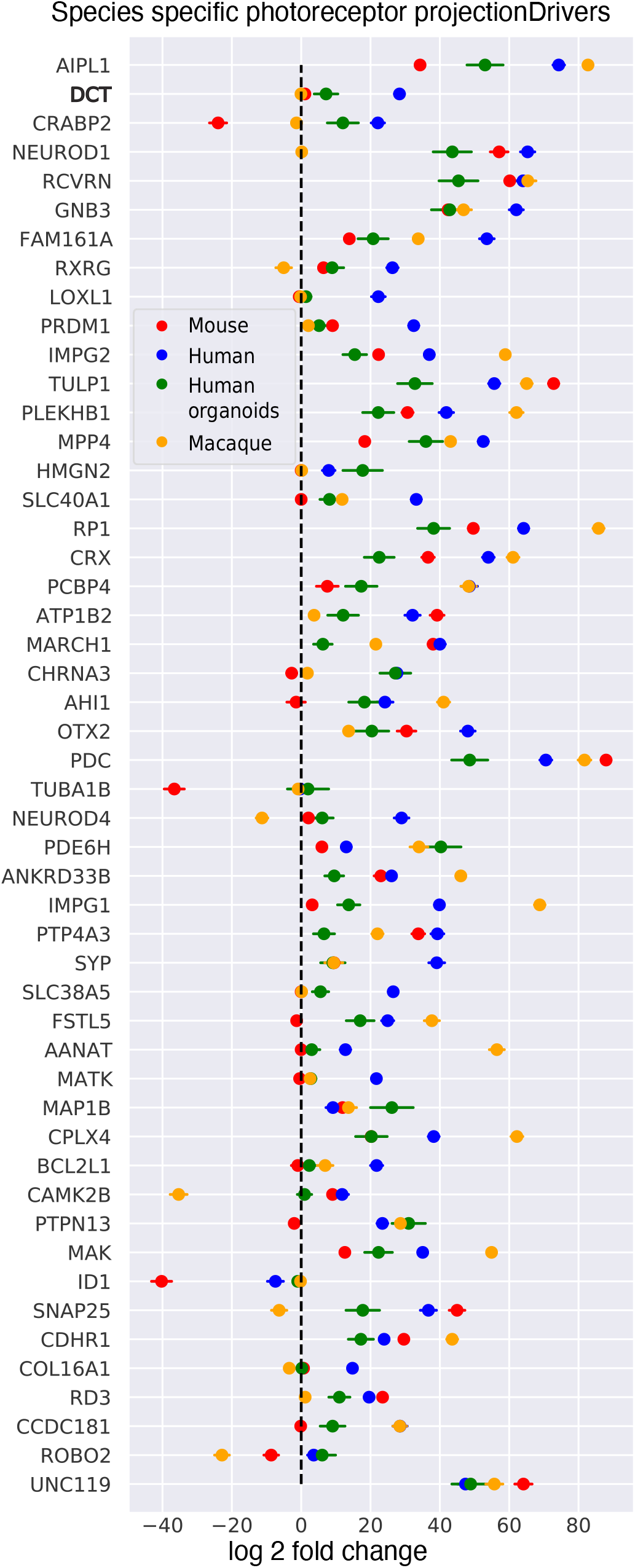
Muti-species comparison of latent space 75 in rods and cones versus all other retinal cell types. Points represent the log_2_ fold change of mean gene expression between rods and cones versus all in these four datasets. Error bars on the points represent the Bonferroni corrected confidence intervals. Genes are ranked by gene weight in latent space 75.

**Figure 5:**
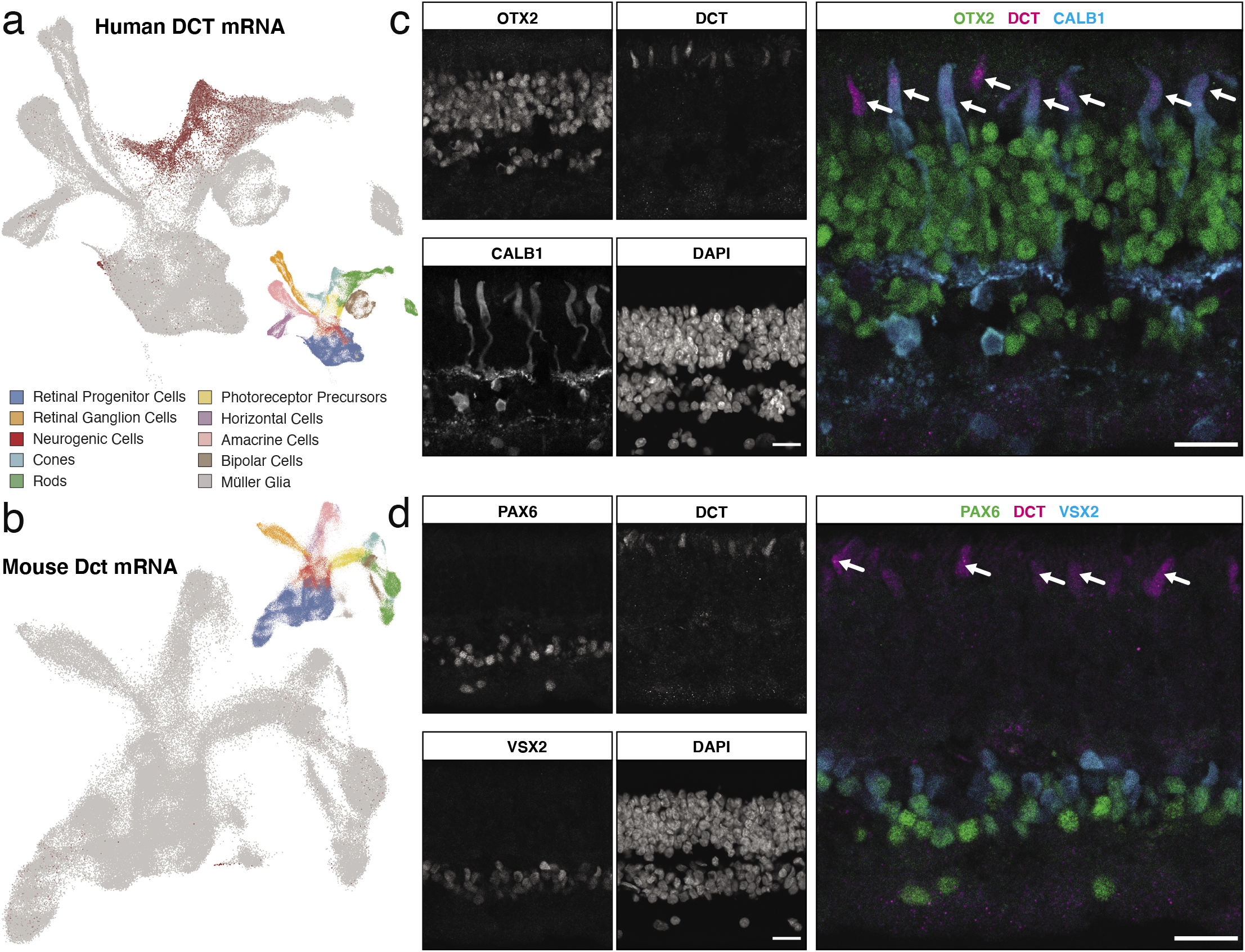
Validation of DCT expression within human cone photoreceptors. A-B) UMAP dimension reduction of the developing A) human and B) mouse retinas, with cells colored by expression levels of DCT. C-D) Immunohistochemistry for DCT (magenta) protein expression in the adult human retina. C) Bipolar cells and photoreceptor nuclei are labeled with OTX2 (green), with cone photoreceptors, horizontal cells, and a subset of amacrine cells labeled by CALB1 (blue). D) Bipolar cells and amacrine cells are labeled by VSX2 and PAX6, respectively. The scale bars indicate 20μm. Arrows in C-D highlight DCT staining within outer segments of cone photoreceptors.

The RPE is known to be intimately involved in both retinal health and function^50,51^ via its role in producing melanin pigment. Recent genetic analyses of ocular albinism identified causal variants within the DCT coding sequence that result in hypopigmentation of the RPE, skin and hair of human patients^52^. Additional features of ocular albinism include bilateral foveal hypoplasia and reduced visual acuity; the cause of which has remained elusive.

Our scProject analyses of the developing human retina demonstrates that human photoreceptors, specifically cones, also robustly express DCT. This is in contrast to mice in which DCT is restricted to the RPE^53^, and displays sparse expression within the mouse scRNA-seq dataset (Fig. 5b). Upon closer examination, we observe enrichment of both *DCT* transcript (Supplement Fig. 4a-c) and DCT protein within human cone photoreceptors (Fig. 5a,c-d); cells that are preferentially concentrated in the fovea and responsible for color vision.

### Scalability of scProject to atlas-level singlecell datasets

While biological conservation of processes within tissues across species is well established, we and others have previously demonstrated that transfer learning can lead to the discovery of conserved biological processes across numerous tissues and cellular contexts in the same species as well^12,14,32^. Thus, to demonstrate the scalability of scProject, we next evaluated our ability to project a large atlas of single-cell gene expression into a reference latent space^54^.

The developing human target dataset consists of 4,062,980 single-cell transcriptional profiles from 112 fetal tissue samples sequenced to a median read depth of 863 UMIs. The dataset was projected into the 80 latent spaces learned in the developing mouse retina. Before projection, we only considered genes shared between the set of latent spaces and the dataset, which left 2403 common genes.

Regression using scProject’s NNLR_ElasticNet took 12 minutes of wall time on one Intel(R) Xeon(R) CPU E7-4809 v4 running at 2.10GHz and took two and half minutes of system CPU time. During this process, scProject has a peak memory usage of approximately 152 Gb. It is worth noting that the TL component is not a contributor to the asymptotic memory usage of an entire analysis as UMAP is far more memory intensive. To address this, we enable users to plot subsets of their dataset using any annotation in the AnnData object. This enhances scalability by allowing users to visualize subsets of large datasets by keying on annotations of interest.

## DISCUSSION

Transfer learning applications have enhanced the utility and application of single-cell gene expression studies, providing mechanisms for the rapid interrogation of biological features across datasets, conditions, and experimental systems. While powerful approaches, they are often restricted to assessing ‘geneset-level’ similarities or differences between datasets, limiting their ability to identify specific genes responsible for differential latent space usage. Here, we describe and validate a new python framework, scProject with projectionDrivers, for evaluating latent space usage across diverse experimental conditions and identifying latent space genes with consistent or divergent expression across projection conditions.

While generalizable to numerous applications, scProject with projectionDrivers is particularly powerful for cross species analysis. Ethical limitations on human subjects and samples require the frequent use of model organisms, yet potentially significant differences in biological processes between species often confound study interpretations. Transfer learning reveals which biological processes are conserved and divergent across species in highly specific contexts. Coupled with projectionDrivers, this method identifies key genes responsible for species specific differences. Our scProject analyses of the developing human retina highlights that human photoreceptors, specifically cones, robustly express DCT— a gene previously only studied in human and mouse retinal pigment epithelium (RPE) as a result of its association with ocular albinism.

The expression of DCT in human cones indicates that a previously unrealized cell type may have clinical relevance to ocular disease pathologies involving mutations in DCT. Specifically, mutations in DCT have been causally linked to hypopigmentation of the RPE, skin and hair of human patients^52^. However, retinal phenotypes are also observed in patients with ocular albinism, including reduced visual acuity and bilateral foveal hypoplasia. Our projectionDriver analysis and validations of cone expression of DCT during development and in adult tissue suggests additional functions of DCT outside of the RPE, warranting further investigations as to the extent to which DCT mutations in ocular albinism patients contribute to visual acuity deficits and foveal hypoplasia.

In addition to highlighting species specific gene usage, we have illustrated the utility of scProject and projectionDrivers in model organism selection. Atlas initiatives are underway to characterize an expanding compendium of human and model organism cell types. Given the scalability of scProject and projectionDrivers as demonstrated by the rapid analysis of over 4 million cells, investigators will be able to quantitatively compare across multiple species using these methods. Thus, scProject and projectionDrivers are powerful tools for optimizing the choice of model organism with which to study a given disease.

## METHODS

### Data Processing and Handling

Single-cell RNA-sequencing files were downloaded through GEO with the following accession numbers: GEO: [GSE11610624](https://www.ncbi.nlm.nih.gov/geo/query/acc.cgi?acc=GSE116106), GEO: [GSE12297024](https://www.ncbi.nlm.nih.gov/geo/query/acc.cgi?acc=GSE122970), GEO: [GSE13800224](https://www.ncbi.nlm.nih.gov/geo/query/acc.cgi?acc=GSE138002), GEO:[GSE11848046](https://www.ncbi.nlm.nih.gov/geo/query/acc.cgi?acc=GSE118480), GEO: GSE14366945 and GEO: GSE15679354. Preprocessing for each dataset was performed as previously described in the relevant publications.

### Code Availability

Each analysis is available as a stand alone jupyter notebook at https://github.com/gofflab/scProjectNotebooks.

### Analysis

Details of the mathematical and statistical methods developed here are provided in Supplementary File 1.

### Acquisition of Human Tissue

Human retinal tissue was obtained from cadaveric eye globes donated for corneal transplant (Mid-America Transplant; St. Louis, MO). Tissue was obtained through a material transfer agreement (MTA) between Mid-America Transplant and the Washington University School of Medicine Department of Ophthalmology and Visual Sciences. Tissue included in these studies was obtained from a 64-year old male, 12 hours post-mortem.

### Tissue Preparations and Immunohistochemistry

Adult peripheral retinal samples were dissected into 1cm x 1cm pieces and fixed for 1 hour in 4% paraformaldehyde in PBS at 4C. Tissue was then washed in 1X PBS, and placed in 30% Sucrose in PBS, overnight at 4C, followed by embedding in O.C.T compound (Fisher HealthCare; cat. #23-730-57). Retinal sections were performed at 15μm using a Leica CM1860 cryostat, collected on SuperFrost Plus slides (Fisher; cat.12-550-15). Tissue was allowed to airdry for >30 minutes, and then stored at −20C until processing for immunohistochemistry.

Slides are removed from the freezer, air-dried for 20 minutes and then rinsed in 1X PBS. Slides are then placed in blocking solution (5% Horse Serum, 0.2% Triton X-100, 0.02% Sodium Azide, 0.1% BSA (powder) in 1X PBS) for 2 hours at room temperature. Samples are incubated in primary antibody overnight at 4C, followed by 3 washes in 1X PBS plus 0.05% Triton X-100 for 5 minutes each wash and incubation of slides in secondary antibody within the blocking solution. Following secondary, slides are washed in 1X PBS plus 0.05% Triton X-100, treated with DAPI (1:3000 in 1X PBS for 5 minutes), rinsed in 1X PBS plus 0.05% Triton X-100, and mounted using Vectashield (Vector Laboratories; catalog # H-1500-10) hardset mounting medium. Fluorescence detection is then performed on a Zeiss LSM 800 confocal microscope. Antibodies used in these studies include Rabbit α-DCT (Proteintech, cat. 13095-1-AP; 1:200), Mouse α-Calbindin-D-28K (Sigma, cat. C9848 clone CB-955; 1:200), Goat α-OTX2 (R&D Systems, cat. AF1979; 1:200), Mouse α-Pax6 (DSHB, cat. PAX6a.a. 1-223; 1:200), Sheep α-Chx10/Vsx2 (Exalpha, cat. X1180P; 1:500), Alexa Fluor 488 Donkey α-Rabbit IgG (Invitrogen, A21206; 1:500), Alexa Fluor 568 Donkey α-Mouse IgG (Invitrogen, A10037; 1:500), and Alexa Fluor 647 Donkey α-Sheep IgG (Invitrogen, A21448; 1:500; used for both Sheep and Goat primary antibodies).

## ACKNOWLEDGEMENTS

This work is supported by NIH grants from BRAIN Initiative in partnership with the National Institute of Neurological Disorders K99NS122085 (GSOB), The National Institute of Aging R01AG066768 and R01AG072305 (LAG), the National Science Foundation IOS-1665692 (LAG), the National Eye Institute R00EY27844 (BSC) and P30 EY002687 (BSC), the Kavli NDS Distinguished Postdoctoral Fellowship (GSOB), the Johns Hopkins Provost Postdoctoral Fellowship (GSOB), and by an unrestricted grant to the Department of Ophthalmology and Visual Sciences at Washington University from Research to Prevent Blindness (BSC).

## SUPPLEMENTAL FIGURE LEGENDS

### Supplemental Figure 1

Graphs indicating projectionDrivers for latent space 39. a) The points represent the log_2_ fold change of mean gene expression between cells expressing latent space 39 and those that do not. The bars on the points represent the Bonferroni corrected confidence intervals. The genes are ranked by the product of the log_2_ fold change and the L1 adjusted gene weights in latent space 39. b) The points represent the log_2_ fold change of mean gene expression multiplied by the L1 adjusted gene weights in latent space 39. The genes are ranked by distance from the y-axis.

### Supplemental Figure 2

a) UMAP dimension reduction of the scRNA-seq dataset for the developing mouse retina. Each point represents a single mouse retina cell colored by their corresponding coefficient for latent space 75. b) UMAP dimension reduction of the scRNA-seq dataset for the developing human retina. Each point represents a single human retina cell colored by their corresponding coefficient for latent space 75. Corresponding dimension reductions of both human and mouse datasets colored by annotated cell type are provided as a reference (middle).

### Supplemental Figure 3

Heatmaps indicating gene expression enrichment across annotated cell types in a) adult human and b) adult mouse scRNA-seq datasets, highlighting GADD45G/Gadd45g expression in human (rods) but not mouse photoreceptors (red boxes). Additional listed genes showcase known markers of the various cell types.

### Supplemental Figure 4

Heatmaps indicating enrichment of DCT transcript expression in cones within a) developing human retina, b) adult macaque retina, and c) developing cone and rod photoreceptors in human induced-pluripotent stem cell-derived retinal organoids. Additional listed genes showcase known markers of the various cell types.

## Notes

https://github.com/gofflab/scProjectNotebooks

https://pypi.org/project/scProject/

### Competing Interest Statement

The authors have declared no competing interest.

### Summary of Updates

Added missing Supplemental File containing the explanatory math for scProject and projectionDrivers

