## Supplementary figures and images for "Identifying Gene-wise Differences in Latent Space Projections Across Cell Types and Species in Single Cell Data using scProject"

### Supplemental Figure 1

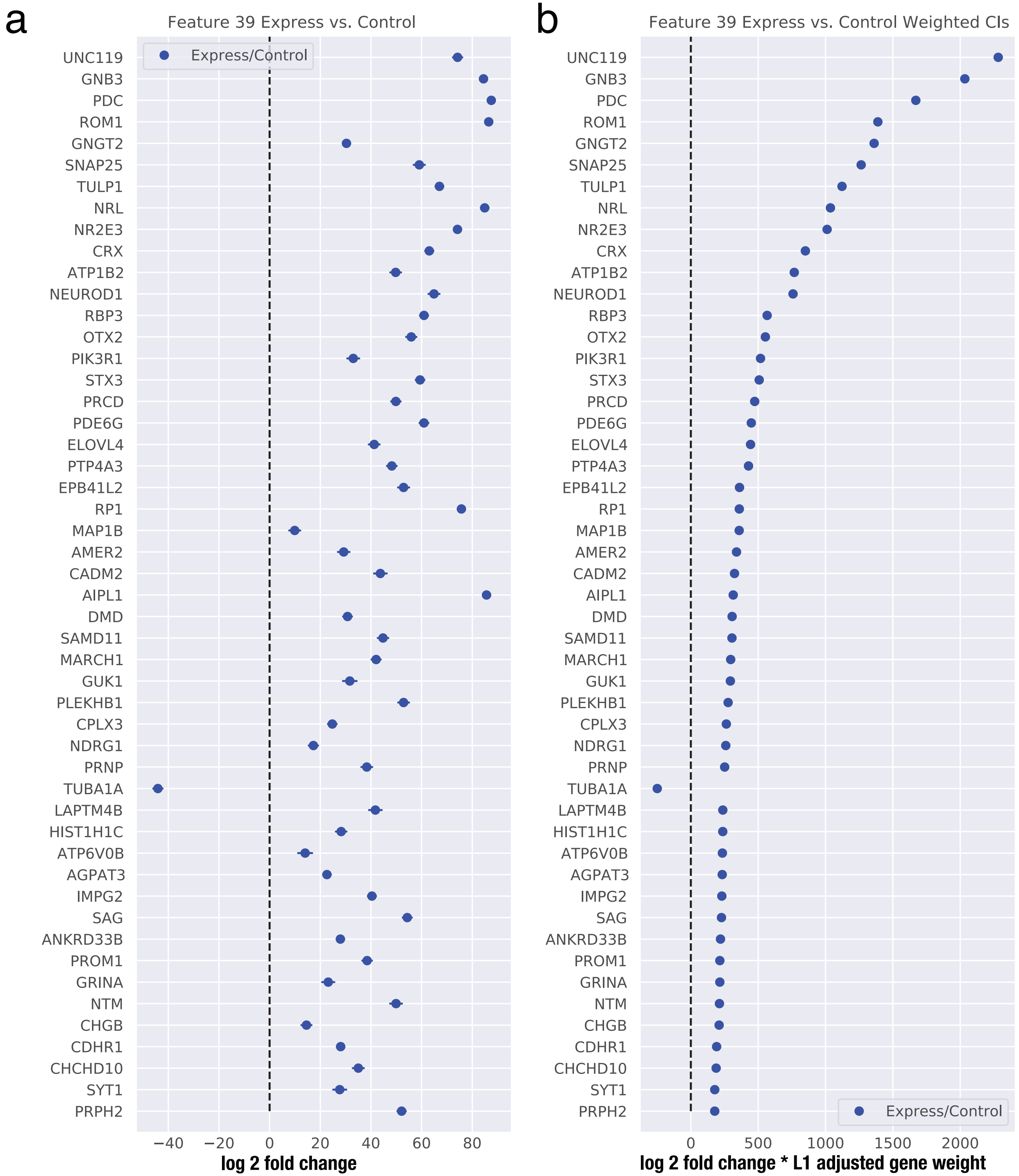

### Supplemental Figure 2

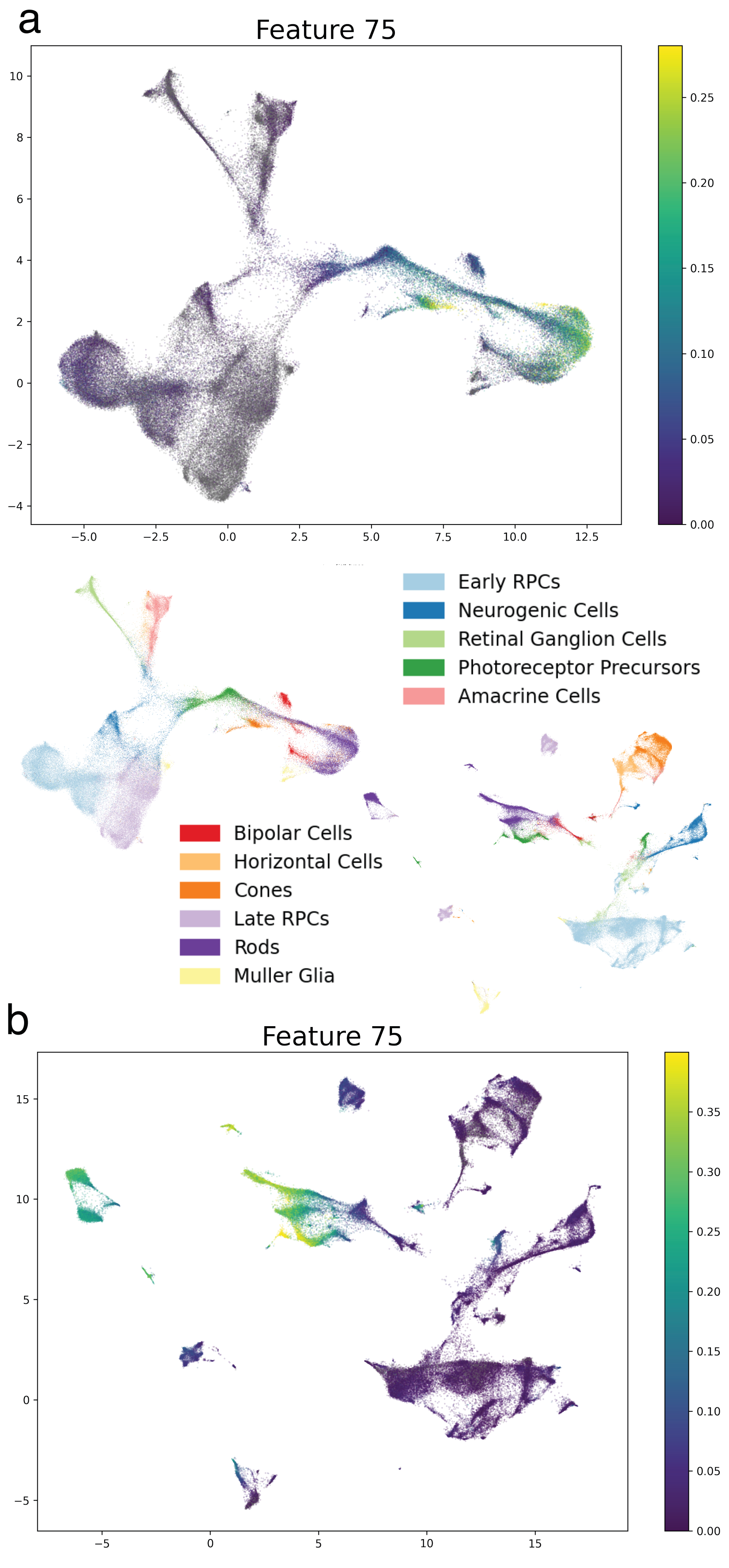

### Supplemental Figure 3

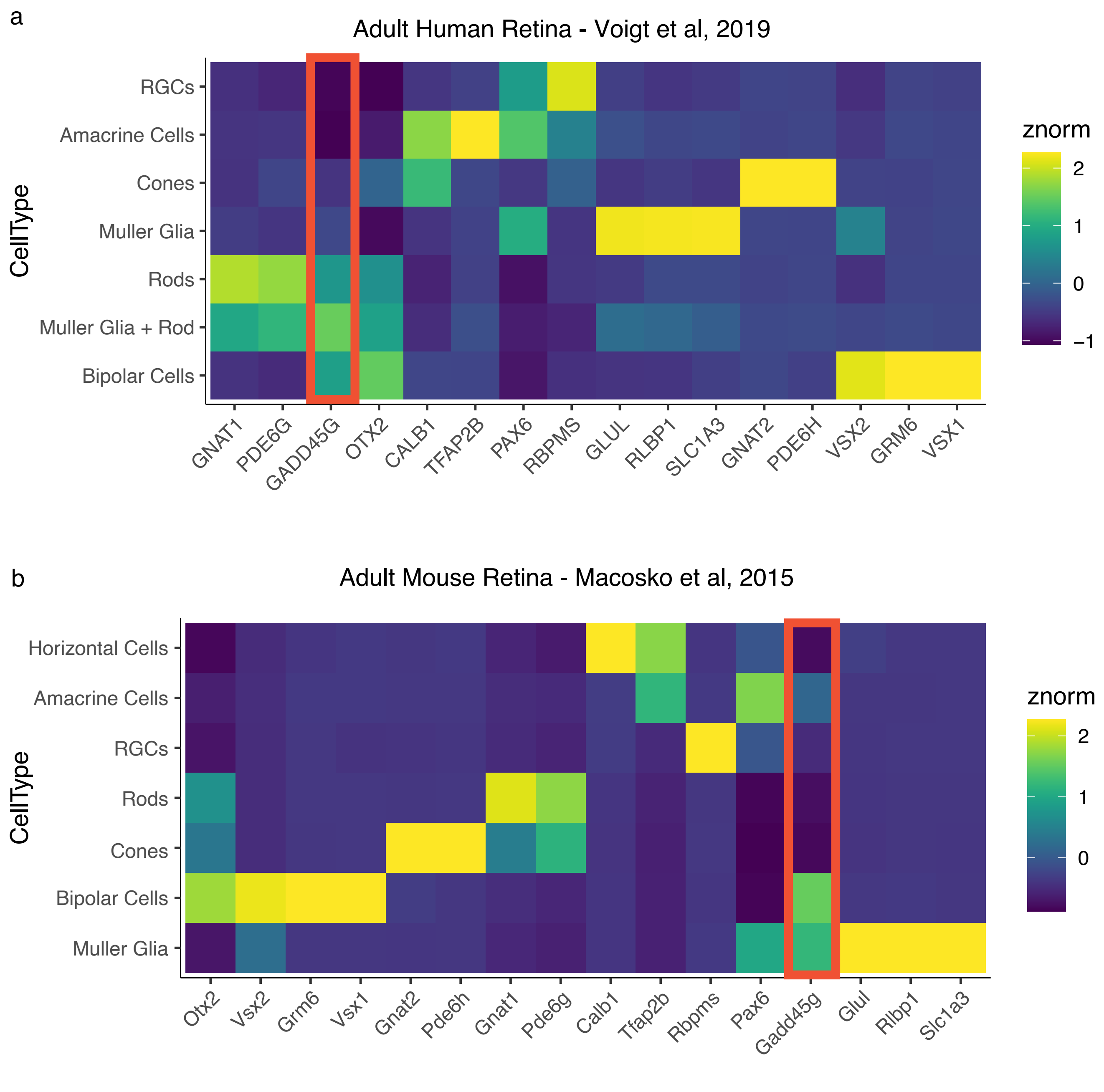

### Supplemental Figure 4

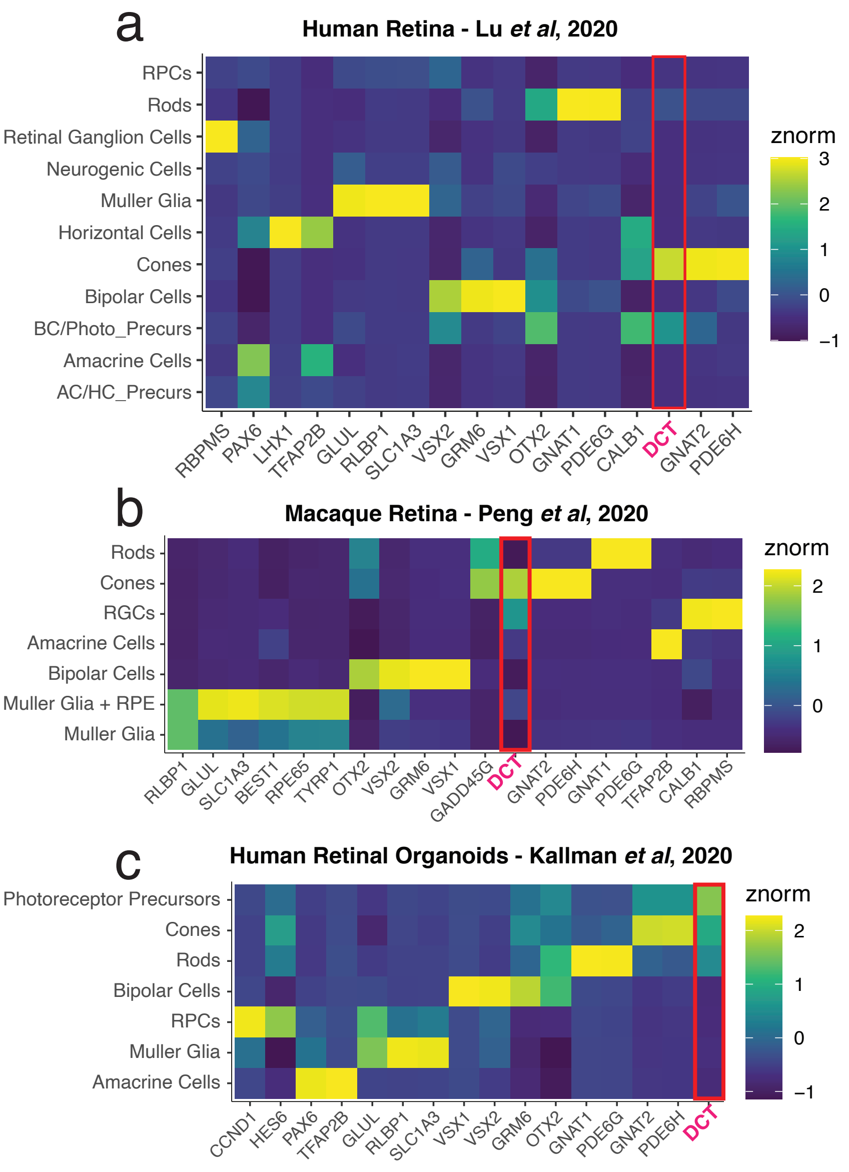
