## Supplemental File 1 for "Identifying Gene-wise Differences in Latent Space Projections Across Cell Types and Species in Single Cell Data using scProject"

### 1 Projection Drivers Statistic:

Overview:
We designed Projection Drivers as a way to quantify which genes in a feature are driving the differential feature expression. Before Projection Drivers, biologists had no rigorous, systematic way to pick out the differential gene expression driving differential feature expression. Simply, Projection Drivers seeks to pick out genes that are both highly expressed in a feature and deferentially expressed between the two clusters of interest. More broadly, Projection Drivers has utility in any feature and loading regression model where it is desirable to pick out attributes of a feature driving differential feature expression.

Let $\hat{\mu_1}$ be the vector of mean gene expression for cluster 1
Let $\hat{\mu_2}$ be the vector of mean gene expression for cluster 2
Let $\rho$ be the dimensionality i.e. the number of genes
Let $n_1$ be the number of observations in cluster 1
Let $n_2$ be the number of observations in cluster 2
Let $A_i$ be the ith column in the amplitude matrix (ith feature/pattern)
Let $\alpha$ be the level of confidence for the intervals
Let $S_j^2$ be the sample variance of gene j

Fix an i(a feature). Divide $\hat{A_i}$ by the number of entries s.t. $\hat{A_{ij}} > 0$. This normalizes the L1 norm of $\hat{A_{ij}}$ to the number of nonzero entries in $\hat{A_i}$.
We define the Projection Drivers confidence intervals:

$$(\hat{\mu_1} - \hat{\mu_2}) \cdot \hat{A_i} \pm t_{n_1+n_2-2,\frac{\alpha}{2\rho}} \sqrt{S_j^2(\frac{1}{n_1} + \frac{1}{n_2})}$$

We define the Bonferroni Corrected Confidence intervals:

$$(\hat{\mu_1} - \hat{\mu_2}) \pm t_{n_1+n_2-2,\frac{\alpha}{2\rho}} \sqrt{S_j^2(\frac{1}{n_1} + \frac{1}{n_2})}$$

Let P be the set of genes whose Projection Driver confidence interval as defined above does not include 0.
Let B be the set of genes who Bonferroni corrected confidence interval as defined above does not include 0.
We classify the set $P \cap B$ as Projection Driver genes.

### 2 Feature Expression Significance:

Overview:
We designed featureExpressionSig as a simple statistical test to test the null hypothesis that the mean expression of a feature in a cluster of interest is less than $\mu$. If we reject the null hypothesis, we can say that the cluster expresses a feature.

Let $\mu$ be the hypothesized expression of the feature
Let M be the mean expression of the feature in question in the cluster
Let $\sigma$ be the standard deviation of the feature expression in the cluster
Let n be the number of samples in the cluster
Let $\alpha$ be the confidence for the t-statistic

We define $T = \frac{M-\mu}{\sigma/\sqrt{n}}$

If $T > t_{n-1,\alpha}$, then we can reject the null hypothesis and say that M is greater than $\mu$ with confidence $\alpha$.

### 3 Hotelling T Squared Test:

Overview:
The Hotelling T Squared test is a multivariate two sample mean test. We evaluate whether the multivariate means of two clusters are significantly different. In term of scRNA-seq data, this test can be used to validate that two clusters of cells that appear distinct using a dimensionality reduction tool are in fact significantly different.

Let $n_1$ be the number of samples in cluster 1
Let $n_2$ be the number of samples in cluster 2
Let $\rho$ be the dimensionality i.e. the number of genes
Let $\hat{\mu_1}$ be the vector of mean gene expression for cluster 1
Let $\hat{\mu_2}$ be the vector of mean gene expression for cluster 2
Let $S_1^2$ be the sample variance covariance matrix for cluster 1
Let $S_2^2$ be the sample variance covariance matrix for cluster 2
We define the pooled sample variance covariance matrix as $S_p^2 = \frac{(n_1-1)\cdot S_1^2+(n_2-1)\cdot S_2^2}{n_1+n_2-2}$
We find $(S_p^2)^{-1}$ using the Moore-Penrose method for a pseudo-inverse of a matrix

We define $T^2 = (\hat{\mu_1} - \hat{\mu_2})(S_p^2)^{-1}(\hat{\mu_1} - \hat{\mu_2})^T$

We define the Fvalue= $\frac{(n_1+n_2-\rho-1)}{\rho(n_1+n_2-2)}T^2$

If Fvalue $> F_{\rho,n_1+n_2-\rho-1,\alpha}$ then we can say that $\hat{\mu_1}$ and $\hat{\mu_2}$ are significantly different at confidence $\alpha$. The implementation in scProject returns a p-value instead of taking in an $\alpha$ value.
